# Extended blood stage sensitivity profiles of *Plasmodium vivax* to doxycycline and tafenoquine using *Plasmodium cynomolgi* as a model

**DOI:** 10.1101/2024.02.23.581752

**Authors:** Peter Christensen, Rosy Cinzah, Rossarin Suwanarusk, Adeline Chiew Yen Chua, Osamu Kaneko, Dennis E. Kyle, Htin Lin Aung, Jessica Matheson, Pablo Bifani, Laurent Rénia, Gregory M. Cook, Georges Snounou, Bruce Russell

## Abstract

Testing *Plasmodium vivax* antimicrobial sensitivity is limited to *ex vivo* schizont maturation assays, which precludes determining the IC_50s_ of delayed action antimalarials such as doxycycline. Using *Plasmodium cynomolgi* as a model for *P. vivax*, we determined the physiologically significant delayed death effect induced by doxycycline (IC_50(96h)_, 1401 ± 607 nM). As expected, IC_50(96 h)_ to chloroquine (20.4 nM), piperaquine (12.6 µM) and tafenoquine (1424 nM) were not affected by extended exposure.

---

The second most prevalent human malaria species, *Plasmodium vivax*, is intractable to continuous growth *in vitro* (1), and consequently, little data is available on the interaction between or on the mode of action of drugs suitable to treat vivax malaria (2, 3). Apart from *in vivo* primate studies, *ex vivo* schizont maturation assays (SMA) remain the only method for investigating *P. vivax* antimalarial drug sensitivity (Russell IJP 2013). However, SMA can only assess the drug’s effect over a single growth cycle (< 48 hours), precluding any investigations on antimalarials with delayed activity such as doxycycline (4). The use of *P. falciparum* as a surrogate for *P. vivax* sensitivity is unsuitable since these two genetically distinct species often differ in their susceptibility and resistance mechanisms to drugs (5, 6) (7) (8). By virtue of its near-identical biological characteristics (hypnozoite formation, early gametocytogenesis, Schüffner dots, etc..) and it close genetic relatedness to *P. vivax* (6), the macaque parasite *P. cynomolgi* has long provided an excellent alternative model. A *P. cynomolgi* line (K4) that has been recently adapted to continuous *in vitro* cultivation now offers, for the first time, the possibility to carry out laboratory-based drug investigations akin to those on *P. falciparum* on this vivax-like parasite. We present investigations on clinically relevant IC_50_ sensitivity profiles of various drugs, including doxycycline, which is associated with a delayed parasite death phenomena (4), using this *P. cynomolgi* line.

*P. cynomolgi* K4A7, a clone of K4, was maintained in *Macaca fascicularis* (*Mf*) red blood cells, as previously described (6) with minor modifications: 25 mM HEPES, 400 µM hypoxanthine, and 5 g/L Albumax II and 10% horse serum (Gibco™ cat. no. 26050088) instead of 20% *Mf*. Tightly synchronized ring stage cultures were prepared by purification of schizonts using a magnetic separator (MACS®, Miltenyi Biotec, Germany) (9) followed by sorbitol treatment (10) 3 hours later. The synchronized state was maintained by repeated sorbitol treatments during schizont rupture every 46 - 50 hours.

Doxycycline (Sigma cat. no. D9891), tafenoquine (Sigma cat. no. SML0396), piperaquine (Sigma cat. no. C7874), and chloroquine (Sigma cat. no. C6628) dried at defined concentrations on plates were produced prior to testing as previously described (11) with the following modifications. Chloroquine and doxycycline were serially diluted 14 times in 70% ethanol after an initial dilution in water, in 70% ethanol for tafenoquine, and in 0.05% lactic acid for piperaquine. The plates were then dried before being sealed and stored at 4° C for up to 40 days. Two hundred µL of the synchronized 0.5% parasitaemia young ring stage parasites cultures (2% haematocrit) were added to each well of the prepared drug plates. Each assay was run in duplicate on 3 separate occasions. The plates were incubated in a humidified chamber containing mixed gas at 37° C. The 48-hour assays were conducted as previously described (11), i.e., once a majority of schizonts had developed (38 - 46 hours). Output data was the percentage of developed schizonts after scoring the life stage of 200 parasites in each well. For the 72- and 96-hour assays, plates were removed from the incubator at 72 and 96 hours, respectively. For each well, parasitaemia was measured by flow cytometry using SYBR green (Invitrogen cat. no. S7563, SYBR Green I) and Mitotracker Deep Red (Invitrogen cat. no. M22426) as previously described (12). Parasitaemia was determined using a BD FACSCanto™ II Cell Analyzer after the acquisition of 20,000 events. Percent schizont development or parasitaemia at each concentration were then used to determine IC_50_ by nonlinear regression using ICEstimator version 1.2 (http://www.antimalarial-icestimator.net)(13, 14).

The marginal, though significant, reduction in the mean IC_50_ sensitivity of *P. cynomolgi* to piperaquine and tafenoquine at 72 or 96 hours as compared to that at 48 hours, is likely due to media exhaustion (over the 96h period) or increased parasitaemia and is unlikely to constitute evidence of a delayed death phenotype. Tafenoquine, a pro-drug used to eliminate hypnozoites is thought to exhibit schizontocidal activity independently of the cP450 2D6 metabolizer status (19). We noted that the IC_50s_ of tafenoquine were lower (1.9 – 1.4) µM than those from previous assays of clinical isolates, 2.68 - 16.6 µM (11, 20). This could represent a difference in species response or variation due to cross-resistance present in clinical samples. Tafenoquine is important for the future of *P. vivax* treatment and should be tested further for drug combination effects, particularly considering its long half-life.

Doxycycline clearly induced delayed death in *P. cynomolgi* as has been observed in other species (4), confirming for the first time this phenomenon in a vivax-like parasite. It is interesting to note that the delayed death effect induced by doxycycline is less pronounced in *P. cynomolgi* (IC_50(48hr)_ of 9.1 µM and IC_50(96hr)_ of 1.4 µM) than that recorded for *P. falciparum* (IC_50(48hr)_ of 5.3 µM and IC_50(96hr)_ of 0.45 µM) (4). The basis of the reduced sensitivity of *P. cynomolgi* to doxycycline relative to *P. falciparum* is at present unknown. It will be interesting to ascertain whether this antibiotic also targets the *P. cynomolgi* apicoplast using the fosmidomycin (inhibitor) and isopentenyl pyrophosphate (rescue agent) combination.

The erythrocytic developmental cycle of *Plasmodium* species is complex, with the inhibitory effects of some compounds evident at different growth stages or only at the later generations. Defining these factors for each of the distinct *Plasmodium* species is fundamental to the design of sensitivity assays. The key benefit of 72 and 96 hours extended sensitivity assays is their ability to provide a holistic sensitivity profile, capturing up to two full rounds of schizont maturation, egress, and invasion, as opposed to the single schizont maturation that the SMA allows. Moreover, extended sensitivity assays are less prone to any bias associated with the parasite stage or synchrony. The use of extended *P. cynomolgi* sensitivity assays now make it possible to define the action of antimalarial drugs that more accurately reflects their effect on *P. vivax*. This will also enable mechanistic studies into drug action/resistance and thus provide the ability to fully evaluate novel therapeutics for this important cause of human malaria.

## Acknowledgements

This work was supported by a Japanese Society for the Promotion of Science long-term Fellowship (BR and OK) and core grants to the A*STAR Infectious Diseases Labs (ACYC and LR) from the Agency for Science, Technology and Research (A*STAR), Singapore, and a Start-up University grant from Nanyang Technological University (LR). GS was supported by a grant from the Agence Nationale de la Recherche, France (ANR-17-CE13-0025-01).

---

**Figure 1.**
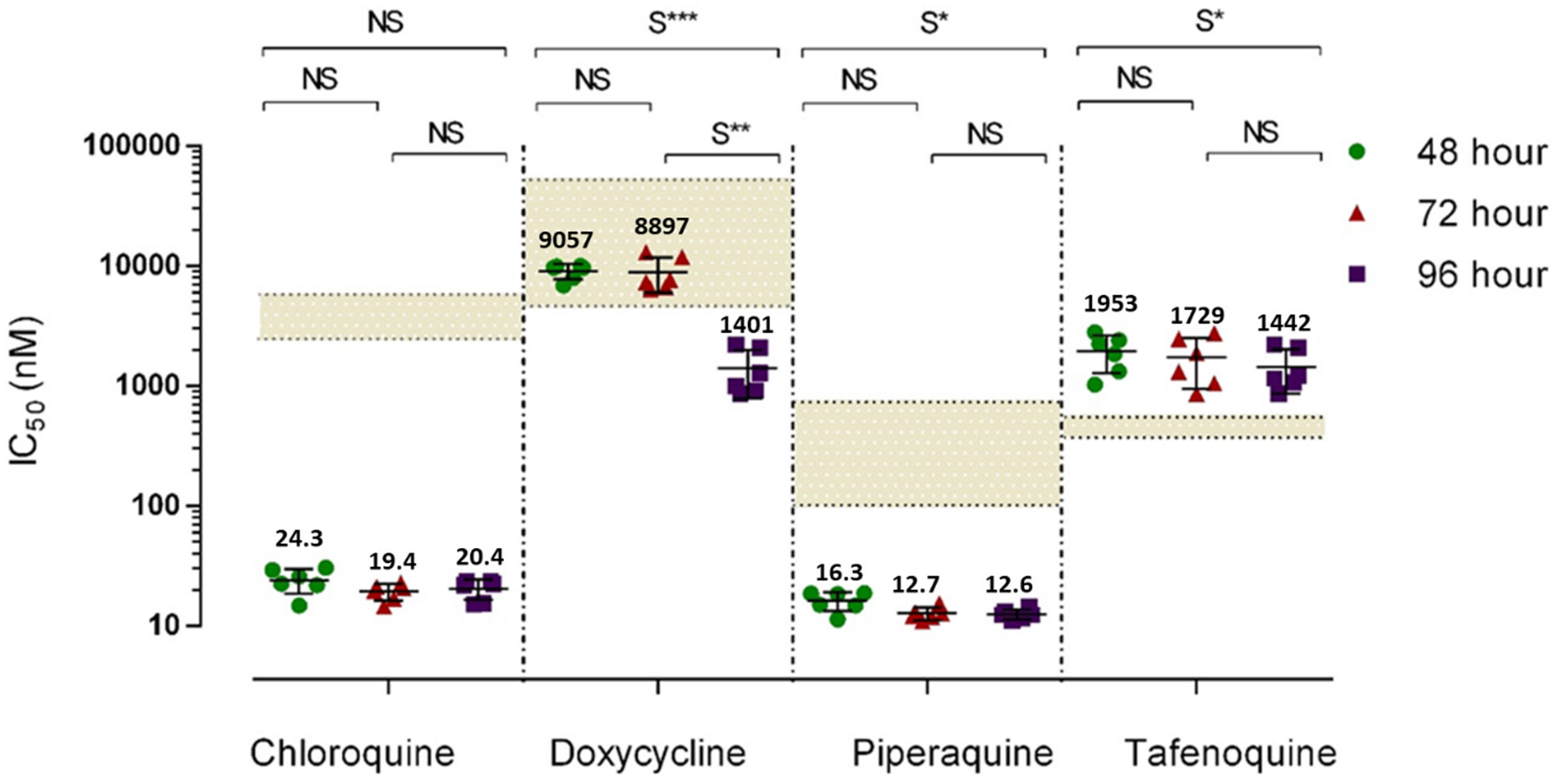
Mean IC_50_ concentrations (nM) of chloroquine, doxycycline, piperaquine and tafenoquine using 48 h, 72 h and 96 h assays. Repeated measures ANOVA with Tukey’s post hoc analysis of significance between assay types are annotated above. Shaded areas indicate previously published *in vivo* plasma C_max_ range for chloroquine (1.25 – 5.08 µM)(15), doxycycline (3.6 – 17.4 µM)(16), piperaquine (79 – 769 nM)(17) and tafenoquine (429 – 481 nM)(18). P value assessment: NS, > 0.05; S*, ≤ 0.05; S**, ≤ 0.01 and S***, ≤ 0.001.

